# Transcriptional regulation of HSCs in Aging and MDS reveals DDIT3 as a Potential Driver of Dyserythropoiesis

**DOI:** 10.1101/2021.09.08.459384

**Authors:** Nerea Berastegui, Marina Ainciburu, Juan P. Romero, Ana Alfonso-Pierola, Céline Philippe, Amaia Vilas, Patxi San Martin, Raquel Ordoñez, Diego Alignani, Sarai Sarvide, Laura Castro, José M. Lamo-Espinosa, Mikel San-Julian, Tamara Jimenez, Félix López, Sandra Muntion, Fermin Sanchez-Guijo, Antonieta Molero, Julia Montoro, Bárbara Tazón, Guillermo Serrano, Aintzane Diaz-Mazkiaran, Mikel Hernaez, Sofía Huerga, Findlay Copley, Ana Rio-Machin, Matthew T. Maurano, María Díez-Campelo, David Valcarcel, Kevin Rouault-Pierre, David Lara-Astiaso, Teresa Ezponda, Felipe Prosper

## Abstract

Myelodysplastic syndromes (MDS) are hematopoietic stem cell (HSC) malignancies characterized by ineffective hematopoiesis, with increased incidence in elderly individuals. In this work, we analyzed the transcriptome of human HSCs purified from young and elderly healthy donors, as well as MDS patients, identifying transcriptional alterations following eight different patterns of expression. While aging-associated lesions seemed to predispose HSCs to myeloid transformation, disease-specific alterations may trigger MDS development. Among MDS-specific lesions, we detected the upregulation of the transcription factor *DDIT3*. Overexpression of *DDIT3* in human healthy HSCs induced an MDS-like transcriptional state, and a delay in erythropoiesis. Such effect was associated with downregulation of transcription factors required for normal erythropoiesis, and with a failure in the activation of their transcriptional programs. Moreover, *DDIT3* knockdown in CD34^+^ cells from MDS patients with anemia was able to restore erythropoiesis. These results identify *DDIT3* as a driver of dyserythropoiesis, and a potential therapeutic target to restore the inefficient erythropoiesis characterizing MDS patients.

**STATEMENT OF SIGNIFICANCE:** This study defines how human aging and MDS development are characterized by transcriptional alterations in HSCs that follow different patterns, some of which may contribute to myeloid transformation. Among them, we demonstrate how MDS-specific upregulation of *DDIT3* in HSCs induces dyserythropoiesis, while its knockdown in HSPCs from MDS patients restores proper erythroid differentiation.

## INTRODUCTION

Hematopoiesis is regulated by different molecular mechanisms that precisely delineate the gene expression programs activated in distinct hematopoietic progenitor cells at specific moments, ultimately enabling the particular functions of each lineage^1^. Deregulation of such mechanisms can lead to the development of various hematological disorders, including myelodysplastic syndromes (MDS), which result from alterations in the first steps of hematopoietic differentiation^2,3^. Thus, MDS are characterized by ineffective hematopoiesis, and clinically manifested as cytopenias and increased risk of transformation to acute myeloid leukemia (AML)^3^.

MDS prevalence is almost exclusive to elderly individuals, with a median age at diagnosis of 71 years and an increased incidence after the sixth decade of life^4^, suggesting that alterations associated with aging predispose hematopoietic progenitors to the disease. In fact, aging is accompanied by a decline in the hematopoietic system that includes defects in both B-cell and T-cell lymphopoiesis, anemia, dysregulation of the innate immune system, and augmented risk of developing myeloid diseases^5,6^. Increasing evidence suggests that the mechanisms intrinsic to HSCs are critical for the adverse hematopoietic consequences seen with age. Some of these mechanisms have started to be elucidated and include changes in the methylome, epigenetic machinery, and histone modification profiles of HSCs^7–10^, along with altered transcriptional programs that include higher expression of genes involved in myeloid differentiation^11,12^. Moreover, in mice, aging causes the expression of genes involved in leukemic transformation in long-term HSCs^11^, while in human, developmental and cancer pathways are epigenetically reprogrammed in elderly CD34^+^ cells^7^, suggesting an aging-mediated predisposition towards the development of myeloid neoplasms. These previous findings suggest that aging-mediated gene expression alterations in HSCs may evolve towards more pathological profiles associated with MDS, although such lesions still need to be elucidated (**Fig. 1A**).

**Figure 1.**
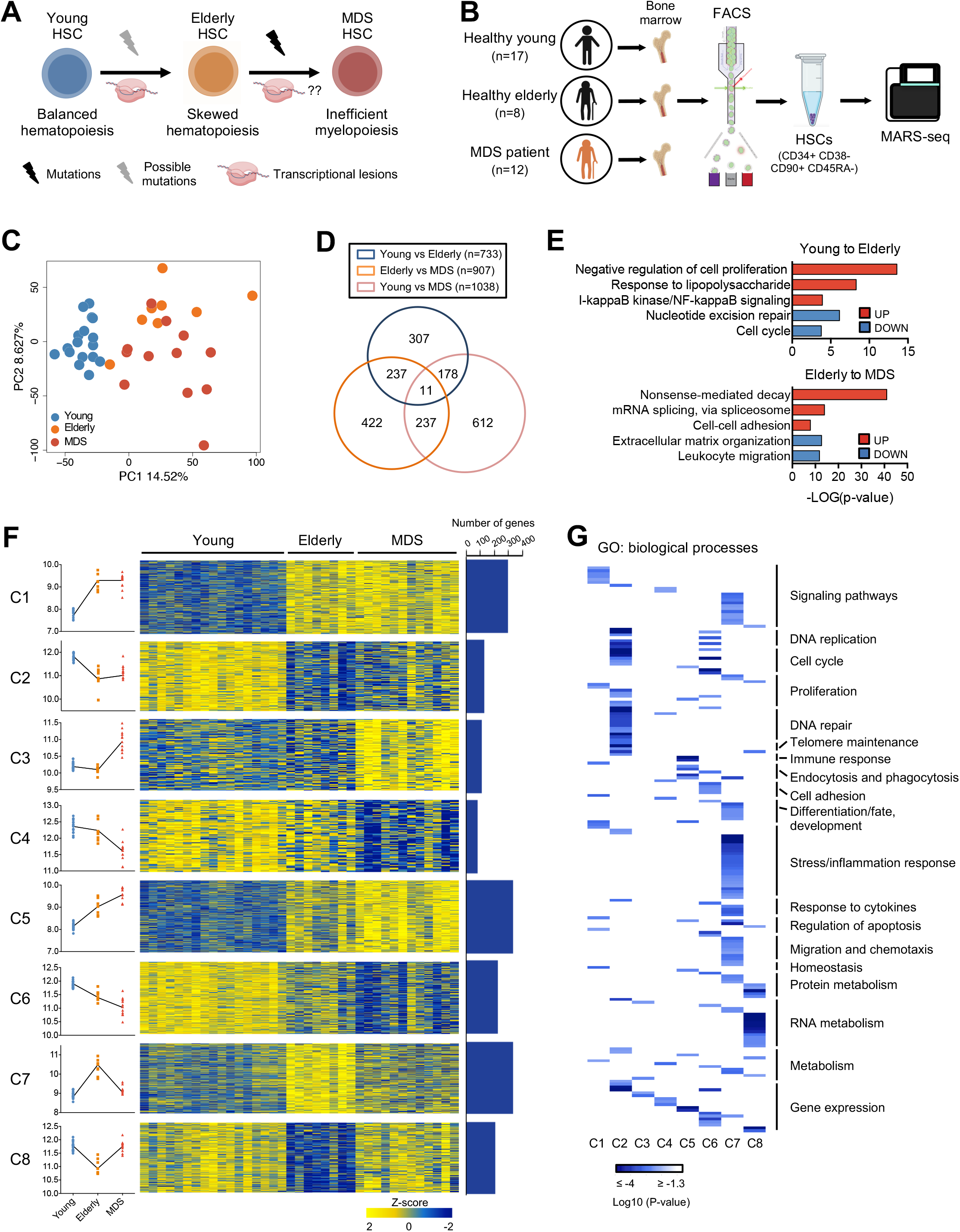
Altered transcriptional profiles of HSCs in aging and MDS. (A) Schematic representation of lesions taking place in aging and in MDS. Aging is characterized by transcriptional lesions of HSCs and can also present mutations in specific genes. MDS HSCs usually present genetic lesions but the transcriptional profile of these cells remains unexplored. (B) Schematic representation of the experimental approach: bone marrow specimens from young and elderly healthy donors as well as from MDS patients were obtained, and HSCs (CD34^+^ CD38^-^ CD90^+^ CD45RA^-^) were isolated using FACS. The transcriptome of these cells was characterized using MARS-seq. (C) Principal component analysis (PCA) of the transcriptome of cells isolated in B. Healthy young (blue), elderly (yellow) and MDS cells (red) are plotted. The percentage of variance explained by PC1 and PC2 principal component is indicated in each axis. (D) Venn diagram representing a partial overlap of genes deregulated in aging (young vs elderly healthy), in the transition to the disease (healthy elderly vs MDS) or between the more distant states (healthy young vs MDS). The number of common or exclusive DEGs is indicated in each area. (E) Barplot showing enriched biological processes determined by GO analysis of genes deregulated in aging or between healthy elderly and MDS HSCs. The -log(p-value) for several statistically significant processes is depicted. (F) Transcriptional dynamisms identified in HSCs in the aging-MDS axis (C1-C8). Left: dot plot depicting the median expression of genes of each cluster in each sample. Each color represents the different states: healthy young: blue, healthy elderly: yellow, and MDS: red. The trend of each cluster is indicated with a line linking the median of each group. Center: heatmap showing z-scores for the expression profile of each cluster of genes in healthy young, elderly and MDS samples. Right: Bar-plot indicating the number of genes per cluster. (G) Heatmap showing the statistical significance (log10p-value) of enrichment of the genes of each cluster in different biological processes, as determined by GO analysis. The different processes have been manually grouped into more general biological functions (right).

So far, many studies aiming to characterize the molecular pathogenesis of MDS have mainly focused on the analysis of mutational profiles associated with the disease, but the fundamental molecular bases of this pathology are still incompletely understood. Recent studies have demonstrated that, as in other hematological malignancies^13,14^, transcriptional mechanisms may also play a relevant role. In this sense, several works aiming to elucidate gene expression aberrations in MDS patients have started to identify, in HSCs-enriched populations, specific lesions with prognostic value or a potential role in the phenotype of the disease^15–29^. Moreover, certain mutations have been associated with specific transcriptional profiles in patients with MDS^17,21–23^, supporting the role gene expression profiles as common integrators of different alterations, being affected by lesions such as mutations or aberrant signaling from the environment, among others.

Although different cell types are phenotypically altered in MDS, previous studies suggest that HSCs are key to understanding this disease due to their privileged position at the apex of the differentiation process^2,3,30–32^. The low abundance of HSCs in the bone marrow makes its characterization very challenging, implying that their expression profile could be partially masked by other cells when HSCs-enriched populations (i.e: CD34^+^) are studied.

In this study, we focused on the analysis of the transcriptional profile of highly purified HSCs from a limited cohort of low-risk MDS patients. By comparing the transcriptional profile of HSCs from healthy young and elderly controls with patients with MDS, we identify specific transcriptional changes associated with aging and with MDS transformation. While aging-associated lesions seem to confer human HSCs with a transformation-prone state, alterations characterizing the progression to MDS may directly alter transcriptional programs leading to inefficient hematopoiesis. Among MDS-specific lesions, we identify *DDIT3,* a transcription factor deregulated in MDS HSCs, as a key regulator of delayed erythropoiesis in MDS. Using gain and loss of function approaches in CD34^+^ cells from healthy donors and MDS patients, we demonstrate the role of this TF in MDS inefficient differentiation indicating that *DDIT3* may be a therapeutic target in patients with MDS.

## RESULTS

### Profiling of HSCs reveals diverse transcriptional dynamics in aging and MDS

FACS-sorted HSCs (CD34^+^ CD38^-^ CD90^+^ CD45RA^-^) obtained from young (n=17) and elderly (n=8) healthy donors, as well as from untreated MDS patients (n=12) were analyzed using low-input RNA-seq (**Fig. 1B and S1A**). Clinical characteristics of MDS patients and controls are shown in Tables S1 and S2. MDS patients included only low- or very low-risk patients with MDS-MLD and MDS-SLD, excluding cases with del(5q), ring sideroblasts or excess blasts, to reduce the heterogeneity associated with MDS patients.

Principal component analysis (PCA) showed that the largest transcriptional differences were observed between young and elderly samples (**Fig. 1C**), with smaller differences between elderly and MDS patients. These observations were confirmed in an unsupervised hierarchical clustering based on the 5% most variable genes among samples (**Fig. S1B**). Differential expression analysis demonstrated 733 genes deregulated between young and elderly healthy samples, and 907 genes between healthy elderly and MDS HSCs (|Fold change (FC)|>2; FDR<0.05) (**Fig. S1C, Table S4**). However, a limited overlap was detected between differentially expressed genes (DEG) in both comparisons (**Fig. 1D**), indicating that specific transcriptional alterations, distinct from those taking place in aging HSCs, occur in MDS. Gene ontology (GO) analyses demonstrated that, among other processes, aging was characterized by an enrichment in inflammatory signaling, and a decrease in cell proliferation and DNA repair signatures; while MDS transcriptional lesions were enriched in RNA metabolism and showed underrepresentation of processes such as migration, or extracellular matrix organization (**Fig. 1E)**. Thus, not only deregulated expression of different genes but also distinct biological processes characterize HSCs in aging and the transition to MDS, suggesting that transcriptional lesions taking place during development of the disease are not a continuous evolution of those found in aging. All together, these data indicate that the transition from healthy elderly to MDS HSCs is characterized by novel transcriptional alterations that lead to deregulation of specific biological processes.

Next, we used MaSigPro^33^ to ascertain the transcriptional dynamics characterizing HSCs in aging and MDS, and identified 8 different patterns of expression: two patterns with genes specifically upregulated or downregulated in aging, (clusters C1, C2); two patterns containing genes with augmented or decreased expression specifically in MDS compared to healthy HSCs (C3, C4); two linear trends harboring genes with either increased or decreased expression in aging and with an exacerbation of such changes in MDS (C5, C6); and finally, two patterns in which genes showed deregulation in aging, and alteration in the opposite direction between healthy elderly and MDS HSCs (C7, C8) (**Fig. 1F**, **Table S5**). Using GO analysis, we identified the main functional categories and their associated biological processes enriched for each cluster (**Fig. 1G, Table S6**). Although some processes such as gene regulation and metabolism were common to different clusters, others were cluster-specific, including DNA replication, cell cycle, telomere maintenance or cell adhesion, among others.

Among other relevant findings, some of which are detailed in the supplemental data (**Fig. S2**), these analyses identified alterations potentially related to MDS development. Processes related to cell proliferation were mainly enriched in C2 but also in C6 (**Fig. S2A, S2E, S2I**), suggesting that an important loss of proliferation activity of HSCs takes place during aging, and it is further exacerbated in MDS, an event that has been previously associated with accumulation of DNA damage^34^. Furthermore, DNA sensing and repair processes were also enriched in C2 and C6 (**Fig. S2A, S2E, S2I**), suggesting a progressive loss of the ability to overcome DNA insults and an increased genomic instability of HSCs during aging and MDS, with a potential direct role in disease development. Interestingly, gene set enrichment analyses (GSEA) demonstrated an enrichment in cancer-related signatures of genes altered in aging, with or without exacerbation of expression in MDS (C1, C2, C5 and C6) (**Fig. 2A**). Furthermore, among these genes, we identified factors with known roles in the development or maintenance of myeloid malignancies (**Fig. 2B**). Some examples included the upregulation of *TRIB1,* a transforming gene for myeloid cells^35^, and the matrix metallopeptidase *MMP2,* which presents high secretion levels in AML^36^; as well as the downregulation of genes such as adenosine deaminase *ADA2,* and the mini-chromosome maintenance proteins *MCM7*^37^ and *MCM4,* whose repression has been involved in myeloid leukemias (**Fig. 2C**). These results suggest that aging-derived transcriptional changes predispose HSCs for transformation.

**Figure 2.**
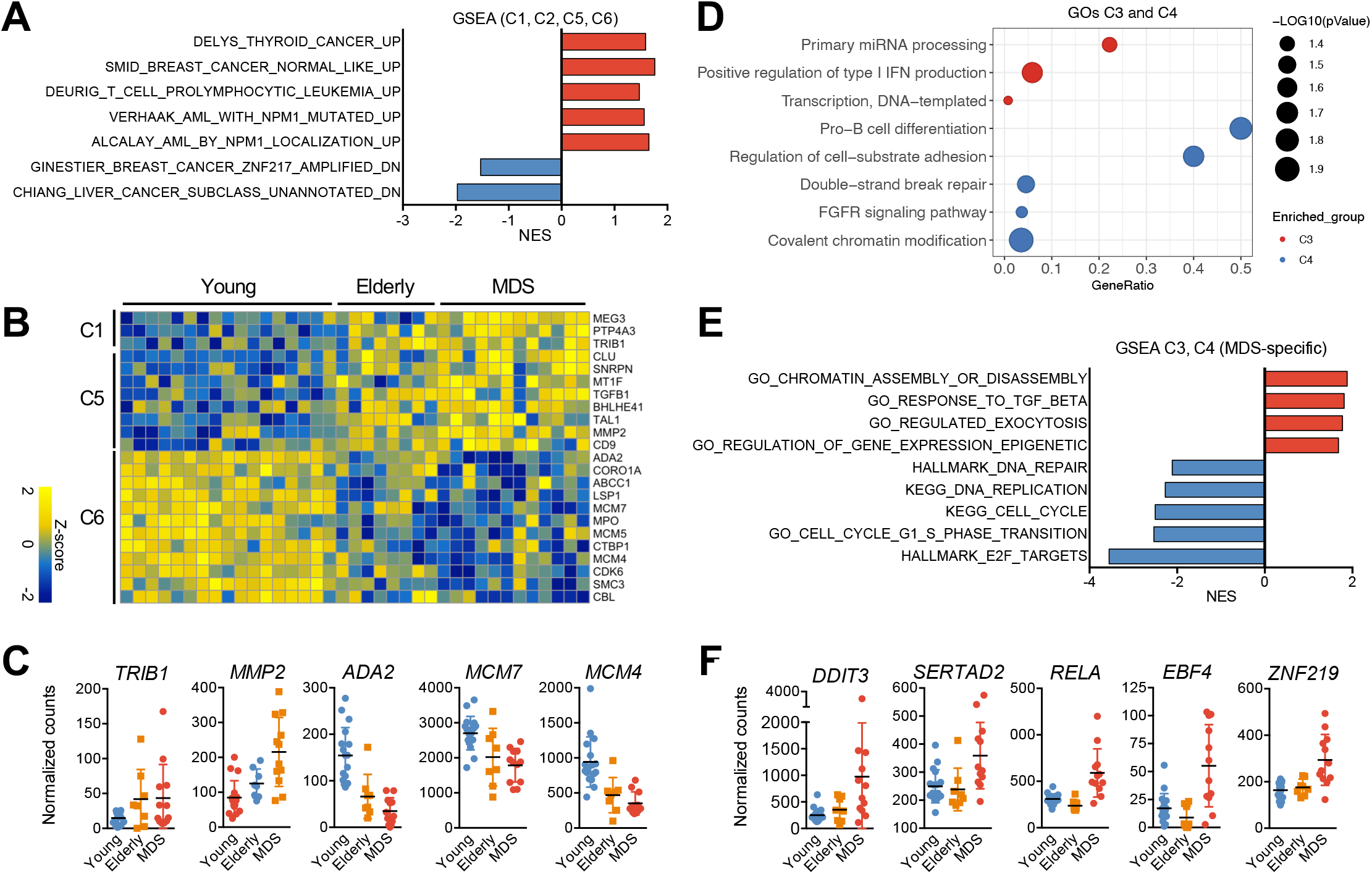
Different transcriptional dynamisms of HSCs in the aging-MDS axis associate with specific biological functions. (A) GSEA analysis of genes from C1, C2, C5 and C6. The normalized enrichment score (NES) for several cancer-related signatures is depicted. Only signatures in which age-matched healthy controls were used as normal counterparts of tumor cells were considered. (B) Heatmap of z-scores of genes with known roles in the development of myeloid malignancies. The cluster to which they correspond, and the gene names are indicated on the left and right side of the heatmap, respectively. (C) Normalized expression of several genes from G in healthy young (blue), healthy elderly (orange) and MDS (red) samples. Each point represents an individual and the mean +/− standard deviation (SD) is depicted. (D) Bubble plot representing statistically significant biological processes and pathways enriched in genes being specifically altered in MDS (C3: red dots; C4: blue dots). Bubble size depicts -log10(p-value) and x axis represents GeneRatio. (E) GSEA analysis of genes from C3 and C4. The NES for several enriched signatures is shown. (F) Normalized expression of several genes from C3 involved in transcriptional regulation in healthy young (blue), healthy elderly (orange) and MDS (red) samples. Each point represents a donor or patient and the mean +/− standard deviation (SD) is shown.

On the other hand, clusters showing exclusive deregulation in MDS (C3, C4) represented lesions with a potential direct role in MDS development. GSEA and GO analyses (**Fig. 2I, 2J**) demonstrated diminished expression of genes controlling cell division and DNA repair (**Fig. S2K**), further suggesting an accumulation of genetic lesions in MDS-associated HSCs. Decreased levels of genes regulating cell-substrate adhesion and B-cell differentiation as well as increased expression of genes involved in processes such as miRNA processing, exocytosis, type I IFN production, or response to TGF-ß were also detected in MDS-associated HSCs (**Fig. S2L**). Finally, increased expression of chromatin and transcriptional regulators, such as *DDIT3, SERTAD2, RELA, EBF4* or *ZNF219,* among others (**Fig. 2F**), suggested an MDS-specific epigenetic and transcriptomic reprogramming that could potentially guide the aberrant phenotype of these cells. Collectively, these transcriptional changes are consistent with an aging-mediated predisposition towards myeloid transformation, and with MDS-specific alterations that may contribute to the development of the disease.

### *DDIT3* overexpression produces an MDS-like transcriptional state and alters erythroid differentiation

To determine the potential functional relevance of the detected transcriptional alterations, we next focused on the transcriptional regulators specifically upregulated in MDS (C3), as they could represent key factors for MDS development. Among them, we selected the TF *DDIT3* (DNA Damage Inducible Transcript *3*/CHOP/C/EBPζ) as one of the top-ranking transcriptional regulators by FC, showing a range of 2.7-10.5 FC over healthy elderly samples in several MDS-derived HSC specimens **(Fig. 2K, S3A)**. Interestingly, increased *DDIT3* expression was only detected in a small percentage of patients when total CD34^+^ cells were analyzed, suggesting its upregulation mainly occurs at the more immature state of differentiation (**Fig. S3B**). In addition, *DDIT3* has been described to be involved in hematopoietic differentiation in mice, specifically promoting abnormal erythroid and myeloid differentiation^38^. To determine the role of its upregulation in HSCs from MDS patients, *DDIT3* was overexpressed in CD34^+^ cells from healthy donors and sorted transduced cells were analyzed by MARS-seq after 2 days in culture in the absence of differentiation stimuli (**Fig. 3A**). *DDIT3*-overexpression induced transcriptional changes with an up- and downregulation of 427 and 128 genes, respectively (|FC|>2, FDR<0.05) (**Fig. 3B**). Consistent with the alterations observed in MDS patients, GSEA demonstrated activation of chromatin remodelers and decrease in DNA repair pathways and cell-substrate adhesion signatures upon *DDIT3* overexpression (**Fig. 3C, S3C, S3D**). In line with the *DDIT3*-induced erythroid bias previously described in mice, we observed an upregulation of genes associated with heme-metabolism or erythroid differentiation, such as *FN3K, GCLM* or *HBB* (**Fig. S3E**). Furthermore, *DDIT3* overexpression promoted an enrichment in cancer-related signatures, indicating a potential oncogenic role of this TF (**Fig. 3D**). More importantly, using our own gene signatures generated from the DEGs detected in our cohort of MDS samples, we observed that *DDIT3*-overexpression induced a “MDS-like” transcriptional state, activating genes upregulated in MDS patients and repressing factors downregulated during the disease (**Fig. 3E-F**). All together, these data suggested that *DDIT3* upregulation plays a key role in inducing a pathological transcriptional state observed in MDS patients.

**Figure 3.**
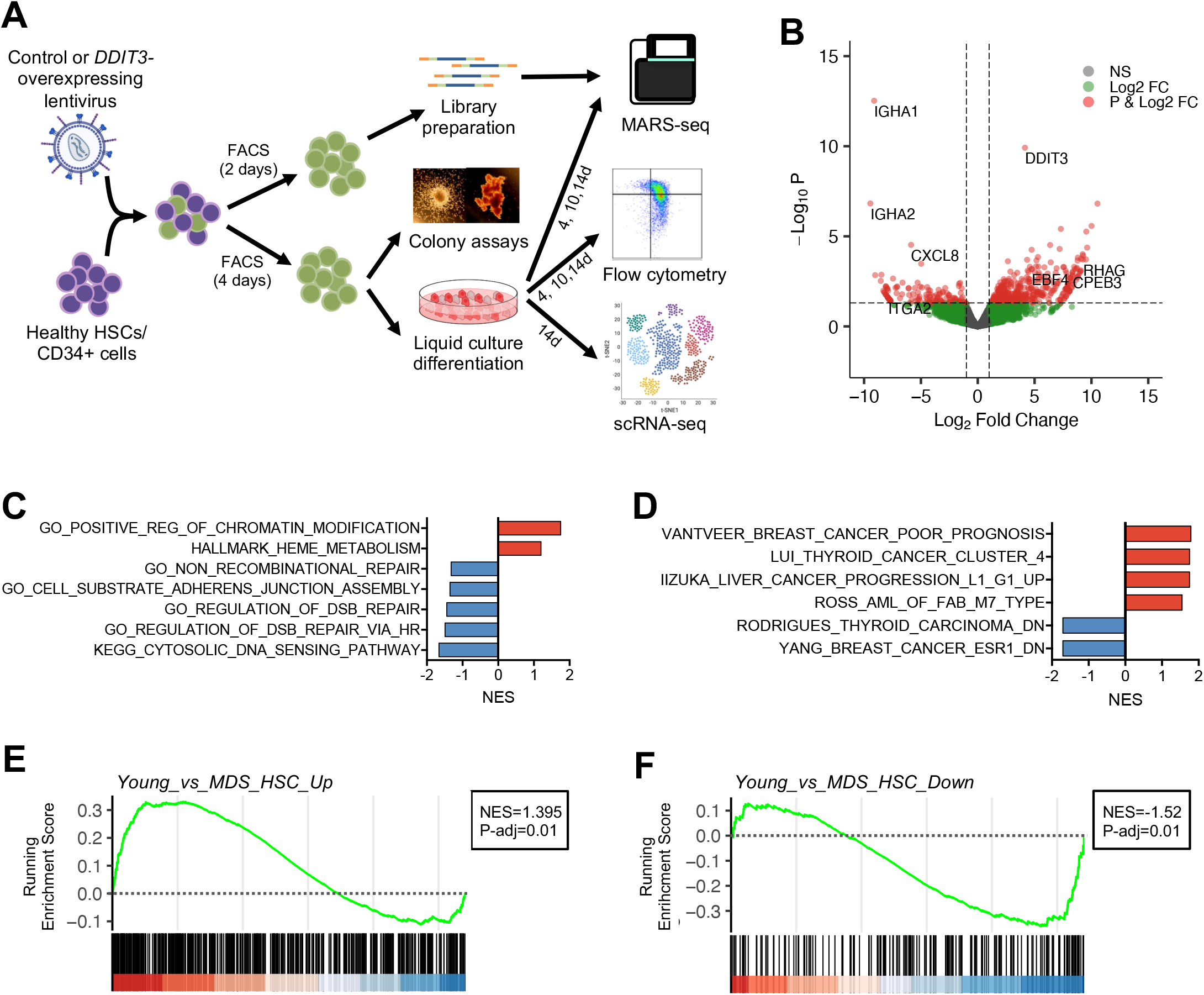
*DDIT3* overexpression promotes and MDS-like transcriptional state. (A) Schematic representation of the experimental procedure: primary HSCs/CD34^+^ cells from healthy young donors were FACS-sorted and infected with a lentiviral plasmid harboring *DDIT3*, or a control plasmid. Two days after infection, transduced cells were sorted and their transcriptome analyzed using MARS-seq. After 4 days of infection, transduced cells were also isolated and used to perform colony and liquid culture differentiation assays. The later were evaluated by flow cytometry and MARS-seq at different time points and by scRNAseq at day 14 of differentiation. (B) Volcano plot of statistical significance (-log10 (p-value)) against fold-change (log2 Fold-change) of gene expression between cells overexpressing *DDIT3* and control HSCs. Green points represent genes with |FC|>2, red points depict genes with |FC|>2 and FDR>0.05 and grey points indicate genes with no relevant changes in expression. Several genes with significant up- and down-regulation are indicated. (D-E) GSEA of cells overexpressing *DDIT3* and control HSCs. The NES for several signatures related to general biological processes (C) and cancer-related signatures (D) are depicted. (E-F) GSEA plots depicting the enrichment upon *DDIT3* overexpression in gene signatures representing genes up- and down-regulated in MDS. The NES and adjusted p-values are indicated for each signature.

We next examined the effect of *DDIT3* overexpression on the function of HSCs using an *ex vivo* myeloid differentiation system starting from primary human healthy CD34^+^ cells (**Fig. 3A**). We observed a statistically significant decrease in the number of both erythroid burst-forming units (BFU-E) and granulo-monocytic colony forming units (CFU-G/M/GM) after *DDIT3* overexpression (**Fig. 4A, S4A**). Liquid culture differentiation assays demonstrated a delay in the differentiation of *DDIT3*-overexpressing cells, with a statistically significant decrease in later stages of erythroid differentiation (stage IV, CD71^-^ CD235a^+^) at days 10 and 14, and an increase in stage II and III erythroid progenitors (CD71^+^ CD235a^-^ and CD71^+^ CD235a^+^ erythroblasts) (**Fig. 4B-C**). No significant alterations were detected in myeloid liquid culture differentiation assays upon *DDIT3* overexpression (**Fig. S4B**).

**Figure 4.**
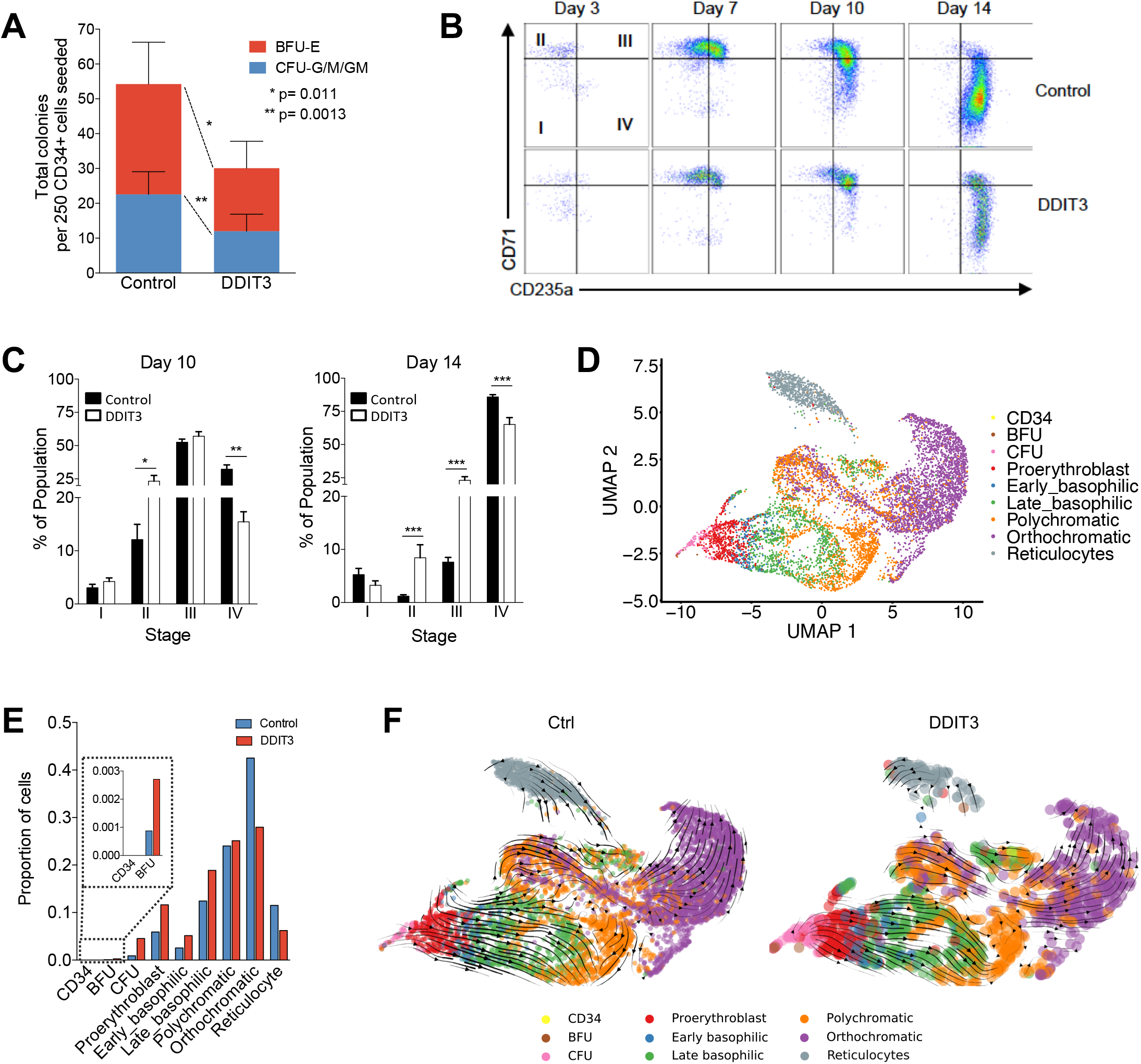
*DDIT3* overexpression promotes a defect in erythroid differentiation. (A) Graph indicating the number of colonies (BFU-E and G/M/GM) obtained for control cells or cells overexpressing *DDIT3*. Statistically significant differences are indicated (*p-value < 0.05; **p-value < 0.01). Biological replicates for this experiment can be found in Figure S4A. (B) Flow cytometry charts representing advanced erythroid differentiation (CD71 and CD235a markers; stages I-IV) for control cells and cell overexpressing *DDIT3,* at the indicated time points. (C) Bar-plots representing the percentage of cells observed in C in stages I-IV at 10 and 14 days of differentiation. Three biological replicates +/− SEM is depicted, and statistically significant differences are indicated (*p-value<0.05; **p-value<0.01; ***p-value>0.001). (D) UMAP plot of the transcriptome of cells subjected to ex vivo differentiation for 14 days. Cells have been clustered in different groups representing erythroid differentiation: CD34^+^ cells, burst forming unit/primitive erythroid progenitor cells (BFU), colony formation unit/later-stage erythroid progenitor cells (CFU), proerythroblast, early-basophilic erythroblast, late-basophilic erythroblast, polychromatic erythroblast, orthochromatic erythroblast, and reticulocyte. (E) Barplot representing the proportions of cells in each of the clusters described in F for control cells or cells overexpressing *DDIT3* upon ex vivo differentiation for 14 days. (F) RNA velocity plotted in UMAP space for control or *DDIT3*-overexpressing cells in clusters defined in E. Streamlines and arrows indicate the location of the estimated future cell state.

To further characterize the effect of *DDIT3* overexpression in erythropoiesis, we performed scRNA-seq in human healthy CD34^+^ cells overexpressing *DDIT3* or a mock control after 14 days in ex vivo liquid culture differentiation (**Fig. 3A**). Clusters representing myeloid differentiation were excluded from further analyses and we focused on clusters associated with erythroid progenitors (**Fig. S4C**). Cluster 4 was annotated as reticulocytes, due to its low counts, high ribosomal RNA content and the expression of genes characterizing this stage, such as genes from the ATG family (**Fig. S4C, S4D**). The remaining clusters were annotated as different stages of erythroid differentiation according to published transcriptomic data^39^ and included CD34^+^ cells, burst forming unit/primitive erythroid progenitor cells (BFU), colony formation unit/later-stage erythroid progenitor cells (CFU), proerythroblasts, and different stages of erythroblasts (early-basophilic, late-basophilic, polychromatic and orthochromatic) (**Fig. 4D**, **S4E**). Expression of *DDIT3* was upregulated in every erythroid differentiation stage in comparison with control cells (**Fig. S4F**), and such overexpression was associated with an increase in the percentage of early erythroid progenitors and a 34% and 46% decrease in the orthochromatic and reticulocyte stages, respectively (**Fig. 4E**). Analysis of differentiation trajectories showed differences in RNA velocity between control and *DDIT3-* overexpressing cells that were consistent with the decrease in late erythroid progenitors (**Fig. 4F**). *DDIT3*-upregulated cells showed decreased RNA velocity from the most undifferentiated cells to the polychromatic state, with shorter, thinner and less dense streamlines being present in these cells as compared to control cells, suggesting an impairment of the correct differentiation to poly- and orthochromatic erythroblasts. Altogether, our data indicate that *DDIT3* overexpression in normal progenitor cells leads to inefficient erythropoiesis.

### *DDIT3*-overexpression leads to a failure in the activation of key erythroid transcriptional programs

To define the molecular mechanisms underlying the erythropoiesis defect induced by *DDIT3* overexpression, we analyzed the transcriptional profile of healthy CD34^+^ cells transduced with this TF and subjected to erythroid differentiation (**Fig. 3A**). GSEA of DEGs (**Fig. S5A**) demonstrated an enrichment in transcriptional signatures of stem and early progenitor cells and decreased features of erythroid differentiated cells, such as oxygen transport or erythrocyte development, upon *DDIT3* overexpression (**Fig. 5A**). Expression of hemoglobin genes and enzymes involved in heme biosynthesis (i.e: *PPOX, FECH, ALAS2, HMBS)* showed increased expression during differentiation in control cells but not in *DDIT3*-transduced cells (**Fig. 5B, S5B**). On the contrary, factors characteristic of stem cells, such as *HOXB* genes, *NDN* or *TNIK,* which were progressively repressed in control cells, showed aberrant high expression in *DDIT3*-overexpressing cells (**Fig. 5B, S5C**). We next used the generated scRNA-seq data to analyze *DDIT3*-induced transcriptional lesions at different stages of erythroid differentiation, from CFU to orthochromatic erythroblast. *DDIT3*-overexpression induced an enrichment in hematopoietic stem and progenitor cell signatures in comparison with control cells (**Fig. 5C**), as well as a decrease in the expression of genes related to heme metabolism or oxygen transport at every stage of differentiation. This was also supported by pseudotime analyses showing early hematopoietic progenitor genes downregulated during normal erythroid differentiation *(SEC61A1, CBFB, WDR18)* to be upregulated in *DDIT3*-overexpressing cells (**Fig. 5D, S5D**); while demonstrating decreased expression of genes involved in erythroid differentiation and heme synthesis *(FAM210B, FOXO3, ATP5IF1,* hemoglobin genes) upon *DDIT3* overexpression (**Fig. 5D, S5E**). These data demonstrate that *DDIT3* overexpression impairs the physiological transcriptional regulation of erythroid differentiation, leading to a failure in the repression of immature genes, and in the activation of factors that are required for proper erythrocyte formation.

**Figure 5.**
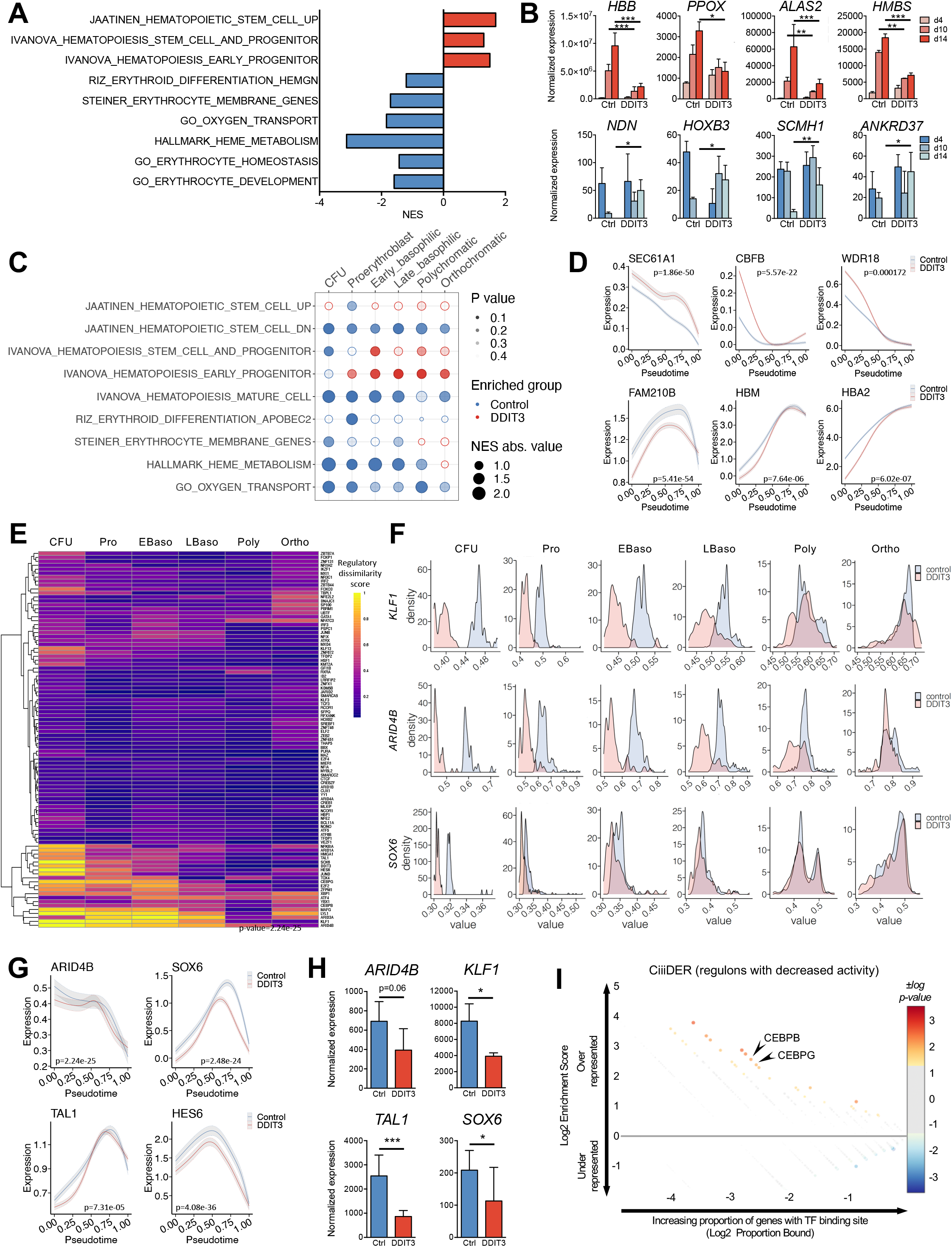
*DDIT3* upregulation leads to a failure in the activation of erythroid differentiation programs. (A) Barplot depicting GSEA analysis upon *DDIT3* overexpression at day 14 of cells overexpressing *DDIT3* and control HSCs at day 14 of erythroid differentiation. The NES for several signatures related to erythropoiesis and stem and progenitor profiles are shown. (B) Normalized expression of erythroid differentiation (top) and stem cell genes (bottom) of CD34^+^ cells at different time points of erythroid differentiation. The mean +/− standard deviation (SD) of 2 biological replicates is depicted. (C) Plot representing GSEA of cells overexpressing *DDIT3* and control cells at different stages of erythroid differentiation detected by scRNAseq. The size of the dots represent NES absolute value, the color indicates the group in which processes are enriched (blue in control cells, red in *DDIT3*-overexpressing cells), and the intensity of the color depicts the p-value obtained for each geneset. (D) Gene expression trends of early hematopoietic progenitor genes (top) and erythroid differentiation factors (bottom) calculated by pseudotime are represented as a smooth fit with the standard deviation of the fit shown in a lighter shade for control (blue) and *DDIT3*-overexpressing cells (red). P-values showing the statistical differences between both trends of expression are indicated. (E) Heatmap showing the regulatory dissimilarity score between control and *DDIT3*-overexpressing cells at day 14 of ex vivo liquid culture differentiation in different stages of erythroid differentiation defined by scRNAseq. (F) Ridge plot showing AUC scores for regulons of *KLF1, ARID4B* and *SOX6* in control (blue) and *DDIT3*-overexpressing cells (pink) at different stages of erythroid differentiation. (G) Gene expression trends calculated by pseudotime of TFs guiding regulons showing decrease activity in *DDIT3*-overexpressing cells, are represented as a smooth fit with the standard deviation of the fit shown in a lighter shade for control (blue) and *DDIT3-* overexpressing cells (red). P-values showing the statistical differences between both trends of expression are indicated. (H) Normalized expression of TFs guiding regulons showing decrease activity in *DDIT3*-overexpressing cells, at different time points of ex vivo erythroid differentiation. The mean +/− standard deviation (SD) of 2 biological replicates is depicted. (I) Analysis of putative transcription factor site enrichment in TFs guiding regulons with decreased activity in *DDIT3-* overexpressing cells using CiiiDER. Color and size of circles reflect p-value of enrichment. Overrepresented transcription factors of potential interest are depicted.

To dwell into the transcriptional programs involved in the abnormal erythroid differentiation induced by *DDIT3* overexpression, we applied a recently described algorithm, SimiC, that infers the differential activity regulons (TFs and their target genes) between two different conditions^40^. Whereas most regulons behaved similarly between control and *DDIT3*-overexpressing cells (**Fig. 5E,** low dissimilarity score, purple), we identified a small number of regulons with differential activity (high dissimilarity score, orange-yellow). Regulons with higher activity in control cells were guided by TFs that positively regulate erythroid differentiation, such as *KLF1, ARID4B, SOX6, TAL1* or *HES6* (**Fig. 5F**), whereas regulons that gained activity upon *DDIT3* overexpression were driven by TFs that negatively impact erythropoiesis, such as *ARID3A, ZFPM1, JUND, ARID1A,* or *HMGA1* (**Fig. S5F**). In most cases, dissimilarities were more evident at earlier stages of differentiation (**Fig. 5F, S5F**), suggesting that aberrant activity of these regulons in immature erythroid progenitors may trigger inefficient erythropoiesis. Differential expression and pseudotime analyses showed that *DDIT3* overexpression was associated with diminished expression of TFs that positively regulate erythropoiesis, such as *SOX6, KLF1, TAL1* or *ARID4B* (**Fig. 5G, 5H**), potentially leading to lower activity of such regulons. Furthermore, the promoters of these TFs were enriched in binding sites for *CEBPB* and *CEBPG* (**Fig. 5I**), which are known to be sequestered and inhibited by *DDIT3*. Thus, these results suggested that abnormally high levels of *DDIT3* in MDS could sequester *CEBPB* and *CEBPG,* leading to decreased expression of their target genes, including TFs with key roles in erythropoiesis, ultimately hampering the activity of transcriptional programs that are necessary for proper terminal erythroid differentiation.

### *DDIT3* knockdown restores erythroid differentiation of primary MDS samples

Based on our results, we hypothesized that inhibition of *DDIT3* in MDS hematopoietic progenitor cells could restore normal erythropoiesis. *DDIT3* was knocked-down in CD34^+^ cells from patients with MDS using shRNAs, and cells were differentiated in liquid culture (**Fig. 6A**). *DDIT3* knockdown in patient 13, a male with MDS-MLD and anemia (hemoglobin (Hb) 11.8 g/dL), led to an increase in the percentage of cells in stage IV (CD235^+^ CD71^-^) at day 7 of differentiation, an effect that was increased at day 13 (**Fig. 6B, 6C**), where cells expressing the control shRNA were partially blocked at stage III. We also detected improved erythroid differentiation upon *DDIT3* knockdown in CD34^+^ cells of four additional MDS cases showing anemia. In two MDS-MLD cases, patient 14 (male, Hb 7.9 g/dL); and patient 15 (male, Hb 9.3 g/dL), *DDIT3* knockdown promoted higher levels of CD71 expression and improved transition to stage III (**Fig. S6A, S6B, S6E**). Furthermore, in the other two cases, one MDS-MLD (patient 16, Hb 11.9 g/dL) and one MDS-EB2 (male, Hb 9.9 g/dL), knockdown of *DDIT3* enhanced the transition to stage IV (**Fig. S6F, S6G**), indicating that inhibition of this factor boosts terminal erythropoiesis. The promotion of erythroid differentiation by *DDIT3* knockdown was validated by transcriptional profiling of cells from two of the patients analyzed (patient 13 and 14) at day 7 of differentiation. This analysis revealed that *DDIT3* knockdown induced increased expression of hemoglobin and heme biosynthetic enzymes, and diminished levels of genes characteristic of immature progenitors, such as HOX genes, compared to cells transduced with a control shRNA (**Fig. 6D, S6C**). Moreover, using published data of gene expression profiling at different erythroid differentiation stages^41^, we observed that *DDIT3* knockdown decreased the expression of genes that are mainly expressed in proerythroblasts, and early and late basophilic erythroblasts, while promoting the activation of genes defining poly- and orthochromatic stages (**Fig. 6E, S6D**). All together, these results suggest that inhibition of *DDIT3* in patients with MDS presenting anemia restores proper terminal erythroid differentiation.

**Figure 6.**
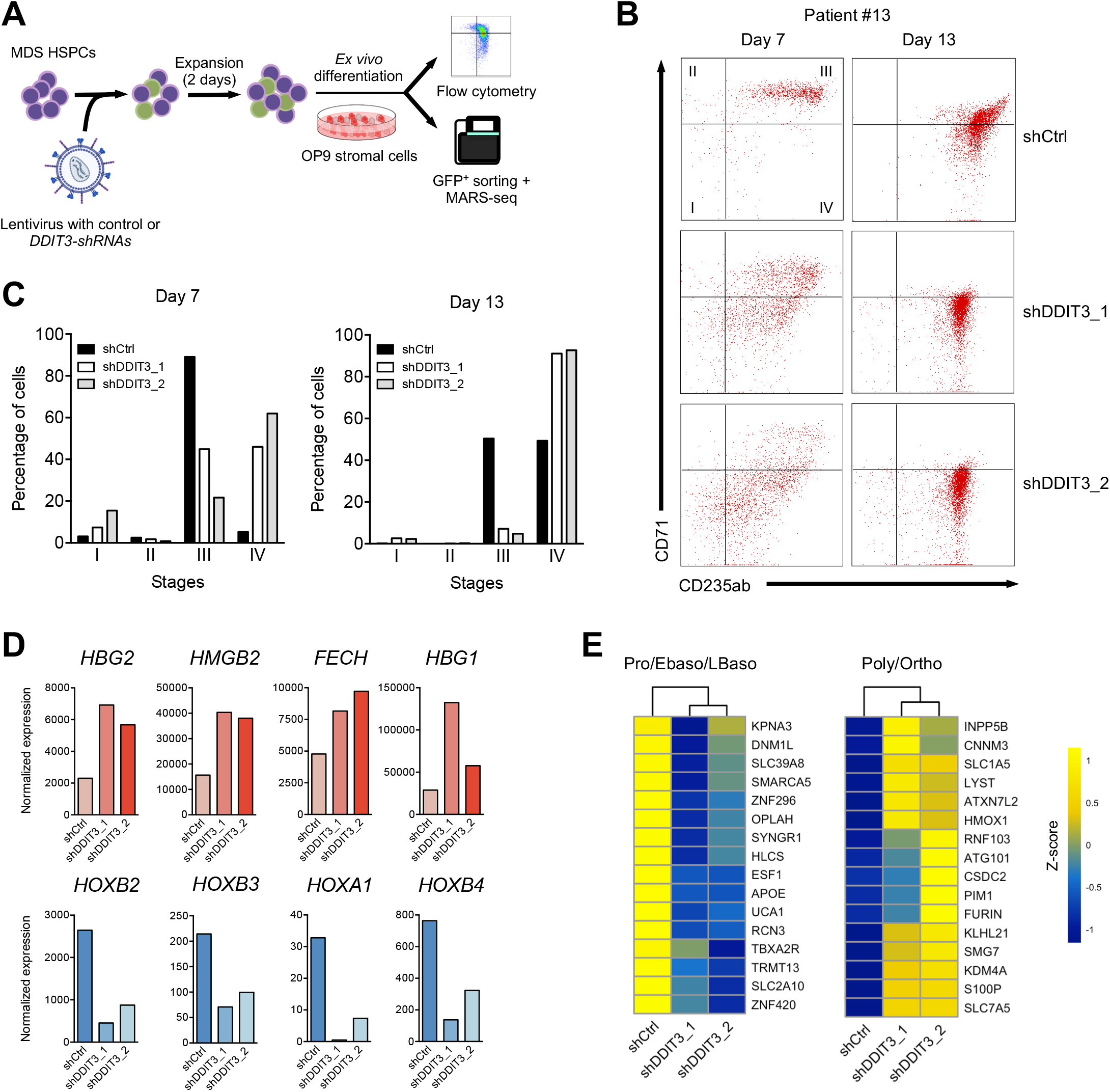
*DDIT3* knockdown in MDS patients with anemia restores erythroid differentiation. (A) Schematic representation of *DDIT3* knockdown experiments in CD34^+^ cells from patients with MDS showing anemia. Cells were transduced with a control or *DDIT3*-targeting shRNA, after two days of infection, cells were subjected to ex vivo liquid culture differentiation over OP9 stromal cells. The differentiation state was evaluated by flow cytometry and MARS-seq analyses. (B) Flow cytometry charts representing advanced erythroid differentiation (CD71 and CD235a markers; stages I-IV) for cells from patient #13 harboring a control shRNA (shCtrl) or shRNAs targeting *DDIT3,* at the indicated time points. (C) Bar-plots representing the percentage of cells observed in B in stages I-IV at 7 and 13 days of differentiation. (D) Normalized expression of hemoglobins (top) and stem cell genes (bottom) of cells transduced with a shRNA control or and shRNA targeting *DDIT3* and subjected to 7 days of ex vivo differentiation. (E) Heatmap of z-scores of genes characteristic of proerythroblasts, early and late basophilic erythroblasts (left), and of genes expressed in poly- and orthochromatic stages (right), for cells transduced with a shRNA control, or an shRNA targeting *DDIT3* and subjected to 7 of ex vivo differentiation.

## DISCUSSION

In this work, we have taken two novel approaches in the study of transcriptional alterations in MDS that we believe are relevant for understanding the molecular pathogenesis of this disease. Firstly, as MDS is considered an HSC disease, we focused our analysis on highly purified HSCs, which may explain the identification of several genes deregulated in MDS not previously described^15–24^. Secondly, we have analyzed the molecular lesions of these cells in the context of aging, allowing for the identification of complex alterations in aging and MDS development that include not only specific lesions of aging and MDS, but also continuous alterations, and even reversal of the aging transcriptome. Our findings suggest a profound transcriptional regulation of HSCs in aging and MDS development and points to the advantage of using age-matched controls for the study of aging-associated diseases.

One of them main characteristics of MDS is the extreme heterogeneity of patients regarding the type and severity of hematopoietic dysfunction, and the molecular lesions that hematopoietic progenitors harbor^42^. In our work, we aimed to partially overcome such heterogeneity by focusing on low or very low-risk MDS-MLD and MDS-SLD, two WHO subtypes that were transcriptionally similar in a preliminary analysis (data not shown), excluding other types of MDS (MDS-RS, MDS-EB, MDS-del(5q), MDS-U). Thus, although the number of cases of study was limited, they represented a relatively homogeneous cohort, which we believe is key in order to find relevant transcriptional alterations. So far, some studies have described an association between genetic and transcriptomic alterations, showing specific gene expression profiles of different FAB and cytogenic groups^22,23^, differential isoform usage of genes in MDS patients carrying mutations in spliceosome and non-spliceosome factors^17^, and alteration of particular subsets of genes with specific splicing mutations^21^. The focus of the present work was not to establish such associations, but to demonstrate how the transcriptomic analysis of purified HSCs can unveil novel transcriptional lesions that may be relevant for MDS development, nevertheless, future works using larger cohorts of HSCs from MDS cases may provide such correlations.

GO analyses unveiled distinct potential functions of the groups of genes identified, undercovering putative transformation mechanisms. Aging-derived lesions suggested a transformation-prone state of elderly HSCs. These alterations included the decreased expression of genes involved in DNA damage sensing and repair pathways, which goes in line with the accumulation of DNA damage observed in human elderly CD34^+^CD38^-^ cells^43^. Diminished levels of such genes might facilitate the accumulation of mutations in these cells, an event that is associated with MDS. Although contradictory data has been published suggesting either increased or diminished proliferation rate of human HSCs, our data support the latest, in line with studies showing that the aging-phenotype of HSCs only emerges after HSCs have reached their maximum number of divisions^44^. Moreover, previous observations showing that DNA repair takes place when HSCs enter the cell cycle^34^ suggest that the diminished proliferative activity of HSCs from elderly individuals could contribute to the accumulation of mutations that takes place in these cells with age^43^. Furthermore, elderly HSCs also showed an enrichment in cancer-related signatures and factors with known roles in the development of myeloid malignancies. Accordingly, murine aged long term-HSCs overexpress genes involved in leukemic transformation^11^, while in humans, a profound epigenetic reprogramming of enhancers has been suggested to promote an aging-driven leukemic transformation-prone state of HSC-enriched populations^7^. Collectively, these data support that aging-dependent alterations in gene expression at the HSC level increase the risk of malignant transformation into myeloid neoplasms, explaining at least in part the higher incidence of these diseases in elderly population.

Our data also identified MDS-associated transcriptional alterations that may trigger the development of the disease, such as upregulation of *DDIT3*, a member of the C/EBP family of TFs, which is involved in functions such as cellular differentiation and proliferation, control of apoptosis and inflammatory processes. *DDIT3* is altered in numerous tumors, with lesions ranging from altered expression (up- and downregulation) to structural abnormalities, including deletions and amplifications (cBioPortal), and translocations that lead to fusion proteins and oncogenic variants^45,46^, suggesting that its role in cancer may be context-specific. Intriguingly, a couple of previous works reported downregulation of *DDIT3*^47,48^ and hypermethylation of its promoter^49^ in MDS. These studies analyzed BM mononuclear cells, from which HSCs represent a very small percentage, and thus, the populations analyzed were different from our work. It is possible that in MDS, *DDIT3* is specifically upregulated in early phases of progenitor cell differentiation and commitment which is consistent with our results, showing that *DDIT3* upregulation was more evident in HSCs than in total CD34^+^ cells. In fact, *DDIT3* has been described as a central regulator of erythro-myeloid lineage specification in mice, where its overexpression, in the absence of pro-differentiative cytokines, enables erythroid programs and impairs myeloid differentiation^38^. Similarly, we observed that, in the absence of differentiation stimuli, *DDIT3* overexpression in human HSCs promoted an erythroid-prone state. Nevertheless, it also promoted a failure in proper erythroid differentiation, with cells being blocked at early maturation stages, indicating that although high *DDIT3* levels seem to prime HSCs for erythropoiesis, they promote an inefficient erythroid differentiation, which is not terminal. Our data suggest that the defect in the activation of differentiation regulons at early stages of erythropoiesis may be due to decreased activity of *DDIT3* binding partners, *CEBPB* and *CEBPG,* which may be sequestered by abnormally high *DDIT3* levels. Future studies characterizing specific subpopulations will determine whether downregulation of this TF takes place at more mature stages, the nature of its binding partners in those cells, and the phenotypic effect that such lesions imply.

The fact that knockdown of *DDIT3* in CD34^+^ cells from MDS cases restores erythroid differentiation, and that it is upregulated in HSC from MDS patients, suggests that *DDIT3* could represent a potential therapeutic target for the treatment of patients, particularly those with anemia. We would like to pinpoint that, besides *DDIT3,* other factors identified in this study showing altered expression in MDS could also have a potential role in the promoting an aberrant differentiation, and their functional role will be evaluated in future studies.

In summary, by analyzing the transcriptome of HSCs we have characterized different transcriptional programs altered during aging and MDS development. Moreover, we have identified *DDIT3* as a key TF involved in the pathogenesis of abnormal erythropoiesis in MDS, thus, representing a potential therapeutic target to restore the inefficient erythroid differentiation of these patients. Similarly, future transcriptional analysis of HSCs in other MDS subtypes may also uncover novel transcriptional lesions with relevant roles in MDS development and progression.

## METHODS

### Sample collection and cell isolation

Bone marrow aspirates were obtained from healthy young (n=17, average=20.53 y/o, range = 18-22 y/o), or elderly donors (n=8, average=67.5 y/o, range=58-81) and untreated MDS patients (n=12, average=70 y/o, range=51-87 y/o) after the study was approved by the local ethics committee, and informed consent was obtained. Sample information and cell isolation strategy can be found in supplemental data and Tables S1, S2 and S3.

### Bulk RNA-seq

Bulk RNA-seq was performed following MARS-seq protocol adapted for bulk RNA-seq, with minor modifications. Exhaustive information about MARS-seq and computational analyses is provided in supplemental methods.

### Mutational analysis

Samples were analyzed either with a custom pan-myeloid panel targeting 56 myeloid genes (CUN), with the panel Oncomine Myeloid Research Assay (ThermoFisher Scientific) (Hospital Val d’Hebron) or with an in-house custom capture-enrichment panel of 92 genes (Hospital Universitario de Salamanca). Details can be found in supplemental data.

### Cell culture

The human 293T cell line and the murine stromal cell line OP9 were cultured as previously described (supplemental methods), and authenticated by short tandem repeat (STR) profiling (Genomics department at CIMAlab).

### *DDIT3* overexpression and knockdown systems

*DDIT3* cDNA was amplified and cloned into the pCDH-MCS-T2A-copGFP-MSCV lentiviral vector. For *DDIT3* knockdown, two different short hairpin RNAs (shRNAs) and a scramble shRNA were cloned into pSIH1-H1-copGFP shRNA lentiviral vector (System Biosciences #SI501A-A). Generation of lentiviruses and further specifications are described in supplemental methods.

### Transcriptome profiling of primary healthy HSCs upon *DDIT3* overexpression

Healthy HSCs were transduced with the overexpression system (details in supplemental methods). Two days after transduction, GFP^+^ cells were FACS-sorted using a BD FACSAria^TM^ IIu in Lysis/Binding Buffer for Dynabeads™ mRNA Purification Kits (Invitrogen), and samples were snap-frozen and kept at −80 C until MARS-seq was performed.

### Colony assays and ex vivo lineage differentiation assays

CD34^+^ cells were incubated in stimulating medium prior to infection with lentiviral particles at multiplicity of infection of 30, as previously described^50^. After 4 days, cells were sorted based on GFP and CD34 (BD Biosciences, #347213) expression on a FACSAria Fusion (BD Biosciences) to perform colony, ex vivo lineage differentiation assays and transcriptomic analyses (details in supplemental methods).

### scRNA-seq of ex-vivo differentiated cells upon *DDIT3* overexpression

Freshly sorted CD34^+^ cells were transduced with lentiviruses as described above and subjected to the ex vivo differentiation system. After 14 days, transduced cells (GFP^+^) were sorted in iced cold PBS1x and 0.05% BSA using a MoFlo Astrios^EQ^ Sorter (Beckman Coulter), and processed according to 10X Genomics Chromium Single Cell 3’ Reagent Guidelines. Sequencing was performed using Illumina NEXTseq500 (Illumina). The procedure and the following computational analyses are explained in supplemental methods.

### Transcription factor binding site analysis

The factors driving regulons with decreased activity in *DDIT3*-overexpressing cells were analyzed using CiiiDER, a TF binding site analysis software. A background gene list was generated by selecting the TFs guiding regulons with low dissimilarity score. Details can be found in supplemental methods.

## Data availability

The data generated in this study are publicly available in Gene Expression Omnibus (GEO) at GSE183328.

## Acknowledgments

This work was supported by the Instituto de Salud Carlos III and co-financed by FEDER funds (PI17/00701, and PI20/01308), CIBERONC (CB16/12/00489); Gobierno de Navarra (AGATA 0011-1411-2020-000010/0011-1411-2020-000011 and DIANA 0011-1411-2017-000028/0011-1411-2017-000029/0011-1411-2017-000030); Fundación La Caixa (GR-NET NORMAL-HIT HR20-00871); and Cancer Research UK [C355/A26819] and FC AECC and AIRC under the Accelerator Award Program. NB was supported by a PhD fellowship from Gobierno de Navarra (0011-0537-2019-000001); MA was supported by a PhD fellowship from Ministerio de Ciencia, Innovación y Universidades (FPU18/05488); TE was supported by an Investigador AECC award from the Fundación AECC, MH was supported by H2020 Marie S. Curie IF Action, European Commission, Grant Agreement No. 898356. We particularly acknowledge the patients and healthy donors for their participation in this study.

## Authorship Contributions

T.E. and F.P. conceived and supervised the study. A.A., J.L, M. S, T.J., F. L., S.M, F.S, A.M., J.M., B.T., S.H, M.D.C., D.V, B.P and J.S.M collected clinical samples and data. A.V., and P.S.M. generated sequencing data. T.E., N.B., C.P., K.R.P. performed ex vivo and in vitro experiments. N.B, M.A, J.P.R, R.O, L.C, G.S, A.D, M.H, A.R, D.F, J.F, M.T.M, and D.L. performed the data analysis. T.E., and F.P. wrote the manuscript.

